# Phylogeny informed international clone assignment with PhyloMLST

**DOI:** 10.64898/2026.07.23.739083

**Authors:** Matthew Neil, Benjamin A. Evans

## Abstract

Multilocus sequence typing (MLST) remains the predominant method for typing bacterial strains. A common method for investigating particularly successful epidemic lineages within a species is to cluster isolates with similar MLST profiles into clonal complexes (CCs). Some CCs, such as international clones (ICs) in *A. baumannii*, are identified with specific sequence types (STs) and are of particular importance to human health. Although theoretically simple, there is a lack of convenient, user-friendly tools perform this analysis. Here we present PhyloMLST, a tool to cluster bacterial isolates into CCs and map them to ICs using the output from existing MLST tools and user-provided STs. As there is potential for aberrant IC assignment arising from excessively large CCs constructed with spurious links, PhyloMLST provides additional functionality to correct IC assignment with a user-provided phylogenetic tree. Although designed with *A. baumannii* in mind, PhyloMLST can be applied to any bacteria where construction and investigation of CCs based on MLST is performed.

**Impact statement:** Many bacterial pathogens are characterised by successful epidemic lineages that are responsible for a substantial number of infections, may be more virulent, and may carry an abundance of antimicrobial resistance genes. These epidemic lineages are comprised of a number of multilocus sequence typing (MLST) sequence types (STs), clustered into clonal complexes (CCs). To date, identifying which STs belong to which epidemic lineage has been challenging, with no simple analytical tools available. Here, we present PhyloMLST - a phylogenetically-aware method for assigning STs to epidemic lineages. The customisable nature of the tool will enable researchers to straightforwardly characterise any population of bacteria that they are working on using MLST data and user-defined definitions of epidemic lineages.

**Data summary:** The PhyloMLST source code and example data shown here is available at https://github.com/Mattn286/PhyloMLST.

## Introduction

Multilocus sequence typing (MLST) is the current standard method for characterising bacteria below the species level. First developed in 1998 (1), the core method involves typing a set of (usually) seven alleles, typically single-copy housekeeping genes. Each allelic variant is an assigned an arbitrary number. The combination of allelic variants, known as an allelic profile, is used to determine the sequence type (ST) of the isolate being analysed. Storing the allele sequence corresponding to each allelic variant and the allleic profiles corresponding to each ST is performed by PubMLST (2).

Isolates of the same ST can be assumed to be very closely related, enabling the characterisation of clades within a species. As a clade grows from a founding member, changes start to accumulate either through mutation or recombination, leading to isolates only differing by a single locus variant (SLV) from the founder, then by a double locus variant (DLV), and so on. To take advantage of this, eBURST was developed, which chains STs separated by SLVs and DLVs into clonal complexes (CCs) (3). The pattern of STs linked toghether within a CC can then be used to infer how the clade expanded from the founder.

As some clades are more successful than others within a species, some CCs are of particular interest, espeiclaly to human health. This has lead to some CCs being given designations. In *A. baumannii*, for instance, globally distributed CCs are known as international clones (ICs). There are currently 12 ICs (IC1-12) (4), with IC2 being the dominant lineage within the species and a serious threat to human health (5).

Despite the straightforward theory behind constructing CCs from MLST data, in practice this analysis can be difficult to perform. While many tools exist to assign isolates to STs, few programs exist to link these together into CCs. PubMLST is available as both a website and through an application programming interface (API), but only returns CC assignments for some species. These assignments are calculated based its entire database, not the user-supplied dataset. An optimised version of the eBURST algorithm known as goeBURST (6) is accessible through PHYLOViZ (7). However, this is complex to install due to a Java dependency and ignores isolates with novel allelic profiles which haven’t been assigned an ST. Neither program is available as a simple tool runnable through a command line interface (CLI) which prevents their use in pipelines, and both are generally difficult to use for large numbers of isolates. This lack of convenient tools to construct CCs has been previously noted as a barrier to analysis of ICs in *A. baumannii* (5), and this limitation likely extends to other species with comparable systems. Perhaps as a result, many studies on *A. baumannii* use well-known STs as a stand-in for the corresponding IC, constricting the size of datasets focusing on specific ICs and limiting analyses to isolates of well-established STs. During the preparation of this manuscript, a Python workflow was published that identifies SLVs and DLVs of a founder ST (8), addressing some of the shortcomings of other tools.

As there is no limit to how many STs can be chained together into a single CC, isolates with very dissimilar allelic profiles can be spuriously linked through many intermediate STs, sometimes described as a “straggly” CC (3). This issue becomes particularly acute when the ratio of recombination to mutation within a species is high, but persists even when this ratio is low. As the number of STs grows rapidly, the formation of spurious links due to the presence of intermediate STs becomes increasingly likely. These spurious links are easy to identify using a phylogenetic tree as isolates within the same CC should cluster closely together. However, to the author’s knowledge, no method has been developed that uses phylogenetic trees to inform CC assignment.

To overcome these issues, we developed PhyloMLST, a CLI tool which can assign isolates to both CCs at a user-defined locus variant level and ICs using user-supplied founder STs. Further, PhyloMLST can optionally use a user-supplied phylogeny to inform IC assignment, correcting the effects of “straggly” CCs. As PhyloMLST was primarily designed for use on *A. baumannii*, we have used IC as a generic term for a named CC. These are equivalent to concepts found in other bacteria, such as *Klebsiella pneumoniae* clonal groups (CGs) (9). Therefore, this tool can be used for any bacteria where MLST is used to construct and assign isolates to CCs.

## Methods

Testing of PhyloMLST was performed using a previously constructed dataset of 1,281 *A. baumannii* chromosomes (10). MLST was performed according to the Pasteur scheme using FastMLST v0.0.16 (11). Founder STs and other STs associated with each IC were obtained from previous literature (4,5,12).

To construct the core genome phylogeny, bakta v1.11 (13) was run on all chromosomes using the light database, and Panaroo v1.5.2 (14) was run using the resulting GFF3 files to produce a core genome alignment with MAFFT (15) as the aligner and core genome threshold of 1. IQtree v2.4.0 (16) was run on this alignment with ModelFinder (17) applied to determine the substitution model. The resulting core genome phylogeny was rooted by an *Acinetobacter nosocomialis* outgroup (GenBank accession no. CP157432.1). Visualisation was performed using TreeViewer (18).

## Results

### Theory and implementation

PhyloMLST is an open-source CLI tool implemented in Python 3. Its only dependencies are pandas (19), BioPython (20), and NetworkX (21), all of which are reliable, widely used, and maintained. PhyloMLST takes as its primary input a TSV containing genome, ST, and marker gene columns. This can be easily generated using FastMLST (11). CCs are constructed at the SLV level by default using the allelic profiles within this table, although the user can choose to cluster profiles with up to six allelic differences. An additional TSV is input to map founder STs to ICs, and CCs containing an isolate of a founder ST are matched to that IC. An optional input can be provided to map any number of additional STs to ICs to match IC assignments to existing literature.

Optionally, the user can supply a phylogenetic tree in Newick format, which PhyloMLST will use to “correct” the IC assignments. This is particularly useful if “straggly” CCs are likely to be a problem. To do this, the tool will search for subtrees which contain below a certain threshold number of isolates that were not assigned to an IC based on locus variant clustering (denoted as ‘Not IC’), and at least one founder ST isolate, then assign all STs in this subtree to the corresponding IC. In this way, only isolates sharing a monophyletic group containing a founder ST and which does not contain above a threshold number of ‘Not IC’ isolates are assigned to an IC, and the rest are demoted to ‘Not IC’. As this correction is done using STs and not isolates, all isolates with numerical STs are guaranteed consistent IC assignment in the final output. The default threshold is zero i.e. a subtree must have no ‘Not IC’ isolates for the STs within to be mapped to the corresponding IC. However, it was found during testing that this is often too stringent due to DLVs (see below). Therefore, the user can optionally raise this threshold.

The final output is a TSV containing isolate allelic profile, ST, CC, assigned IC, and (optionally), corrected IC. The final number of isolates assigned to each IC both before and after correction is printed to stdout,and can be optionally written to a TSV.

### Assignment of *Acinetobacter baumannii* isolates to international clones

PhyloMLST was run on a previously constructed dataset of 1,281 *A. baumannii* chromosomes (10). Initial MLST was performed using FastMLST (11) according to the Pasteur scheme. Clustering of isolates into CCs was performed at the SLV level. Founder STs and other STs previously reported as belonging to specific ICs were obtained from existing literature (4,5,12). These were used to assign isolates to ICs. Phylogenetic correction was performed with a core-genome phylogeny constructed using IQtree v2.4.0 (16), with a subtree Not IC threshold value of 2. Isolates with non-numeric STs (e.g. novel allelic profiles) were masked.

All ICs were mapped to a single clonal complex bar two (Table 1). IC7 was split into two CCs, one containing the founder ST25 and the other containing ST113. ST113 has been previously assigned to IC7 (22), but differs from ST25 by 2 alleles with no intermediate ST linking them in this dataset. IC3 was also split into two CCs, one containing the founder ST3 and the other containing ST229. Similarly to the case in IC7, ST3 and ST229 differ by two alleles with no intermediate ST to link the two in this dataset. ST113 and ST229 were assigned to IC7 and IC3 respectively based on input mapping known STs to ICs rather than CCs. Strains from both these STs were located near to their respective founder ST on the phylogeny, giving strong confidence in these assignments.

**Table 1.** Founding STs, CCs, and total isolates assigned to each IC before and after phylogenetic correction.

| IC | Founding ST | CCs belonging to IC | No. isolates before correction | No. isolates after correction |
| --- | --- | --- | --- | --- |
| IC1 | 1 | 1 | 109 | 109 |
| IC2 | 2 | 2 | 628 | 627 |
| IC3 | 3 | 3, 229 | 39 | 7 |
| IC4 | 15 | 15 | 26 | 26 |
| IC5 | 79 | 79 | 31 | 31 |
| IC6 | 78 | 78 | 27 | 27 |
| IC7 | 25 | 25, 113 | 47 | 47 |
| IC8 | 10 | 10 | 44 | 44 |
| IC9 | 464 | 464 | 9 | 9 |
| IC10 | 33 | 33 | 6 | 6 |
| IC11 | 164 | 164 | 12 | 12 |
| IC12 | 158 | 158 | 5 | 5 |

When assigning IC purely based on CCs, IC3 was unexpectedly large (Table 1). Further analysis found that ST437, corresponding to the ATCC 17978 strain (23), was assigned to IC3. This was surprising as ATCC 17978 is a lab strain and unlikely to be related to recent clinical isolates found in IC3, with ST437 having never been assigned to IC3 to the author’s knowledge. Many of the isolates assigned to IC3 were distributed widely across the phylogeny (Fig. 1), confirming that its large size was due to aberrant clustering at the CC level, not genuine evolutionary trends. Therefore, phylogenetic correction was performed to address this. Due to some ICs containing ‘Not IC’ isolates within their monophyletic groups (Fig. 1), setting a ‘Not IC’ threshold of 0 resulted in loss of likely valid IC isolates. Therefore, the ‘Not IC’ threshold was set at 2. This was sufficient to maintain the coherence IC3 while maintaining other ICs.

**Fig. 1.**
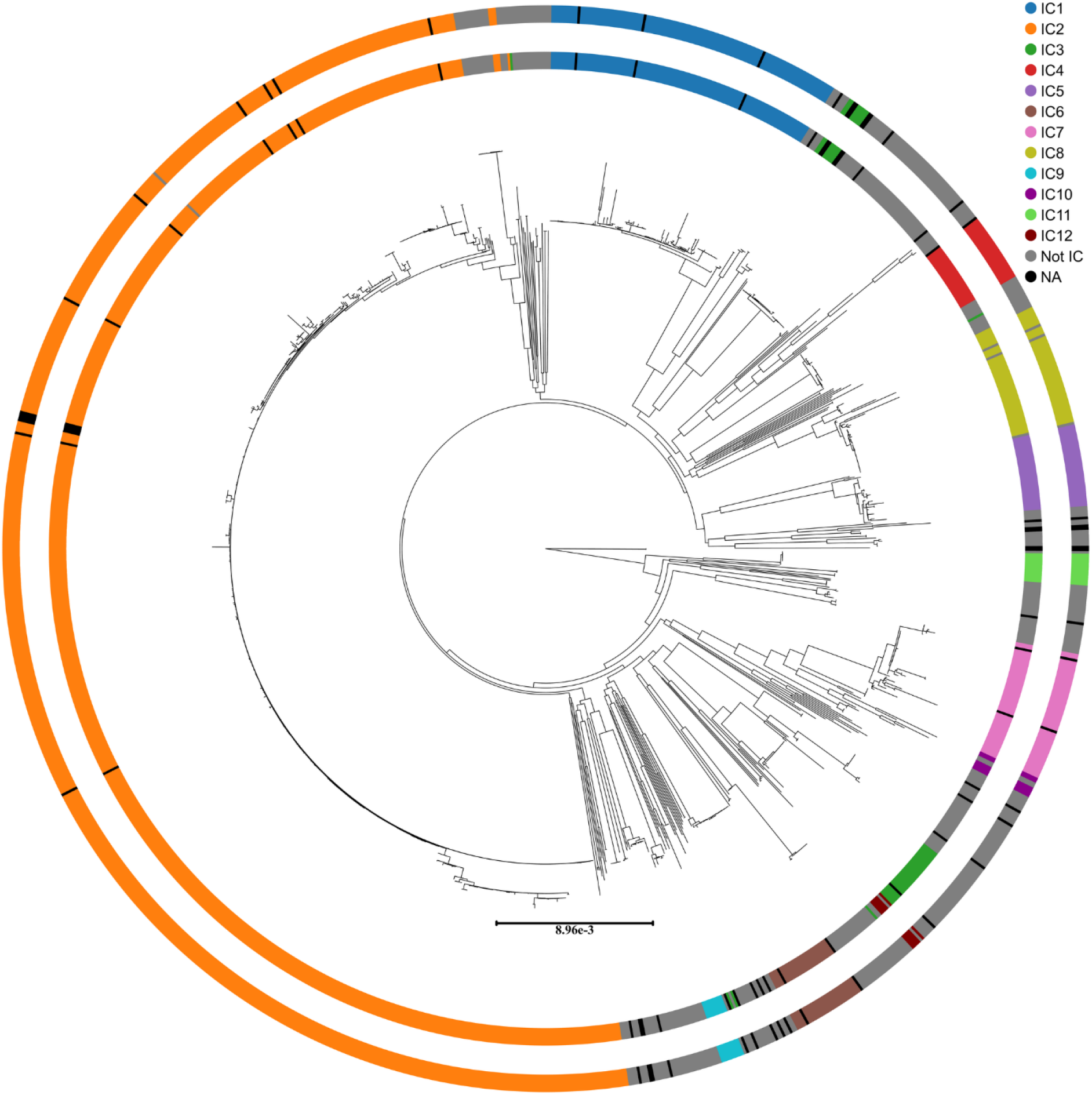
Phylogenetic tree showing IC assignments before and after correction. Phylogeny was constructed from 1,281 *A. baumannii* chromosomes, with the IC of each isolate shown using colored rings as per the legend. The inner ring shows IC assigned using CCs constructed at the SLV level along with predefined STs as described above. The outer ring shows IC after correction using the phylogenetic tree. Scale bar indicates nucleotide substitutions per site. Isolates with non-numeric STs are marked as “NA”.

## Discussion

Although MLST remains the predominant method of typing bacteria, the programs used to cluster isolates beyond the ST level remain lacking. Here, we present PhyloMLST, a user-friendly tool to cluster isolates in CCs and assign them to ICs using user-defined founder STs. To provide maximum control over the output we allow the user to define how many allelic differences to cluster isolates by (SLV, DLV etc.) and input a set of STs which will always map to a corresponding IC. PhyloMLST provides additional functionality by allowing correction of IC assignments using a phylogenetic tree. This can be used to prevent “straggly” CCs causing distantly related isolates to be assigned to the same IC, overcoming the limitations of approaches that construct CCs using ST data only.

We set the default clustering threshold to the SLVs level, and this resulted in some STs that are reported as belonging to the same IC being split into two CCs. Altering the clustering threshold to encompass DLVs would collapse these STs into single CCs. However, we found that using DLVs as the default resulted in spurious clustering of STs that do not belong to the same CC. The phylogenetic correction offered by PhyloMLST overcomes these potential issues to assign isolates to the correct IC.

The main limitation of our approach is that it requires an accurate phylogeny to be constructed from all isolates in a prospective dataset. Recombination can distort phylogenetic trees, potentially impacting the corrected IC assignments. However, recombination mainly distorts branch lengths with limited impact on tree topology (24). As our tool relies on tree topology only, the effects of recombination should be minimal. If recombination is considered to be an issue, numerous tools exist to construct a recombination-free phylogeny for use in correcting IC assignments (25,26).

In conclusion, here we present PhyloMLST, a tool for constructing CCs and assigning ICs based on MLST results. This tool has minimal dependencies and users have a range of options to tailor the output to their needs. To address problems posed by “straggly” CCs, initial IC assignments can be corrected using a user-provided phylogenetic tree, allowing for phylogenetically informed IC assignment. Although we designed PhyloMLST with *A. baumannii* in mind, it can be applied to any bacterial species where MLST is used to cluster isolates into CCs.

## Author Contributions

MN: conceptualisation, methodology, formal analysis, investigation, data curation, writing – original draft preparation, visualisation; BAE: conceptualisation, methodology, resources, data curation, writing – review and editing, supervision, project administration, funding acquisition.

